# Effects of surfaces and macromolecular crowding on bimolecular reaction rates

**DOI:** 10.1101/844753

**Authors:** Steven S. Andrews

## Abstract

Biological cells are complex environments that are densely packed with macromolecules and subdivided by membranes, both of which affect the rates of chemical reactions. It is well known that crowding reduces the volume available to reactants, which increases reaction rates, and also inhibits reactant diffusion, which decreases reaction rates. This work investigates these effects quantitatively using analytical theory and particle-based simulations. A reaction rate equation based on only these two processes turned out to be inconsistent with simulation results. However, accounting for diffusion inhibition by the surfaces of nearby obstacles, which affects access to reactants, led to perfect agreement for reactions near impermeable planar membranes and improved agreement for reactions in crowded spaces. A separate model that quantified reactant occlusion by crowders, and extensions to a thermodynamic “cavity” model proposed by Berezhkovskii and Szabo (*J. Phys. Chem. B* 120:5998, 2016), were comparably successful. These results help elucidate reaction dynamics in confined spaces and improve prediction of in vivo reaction rates from in vitro ones.

## 1. Introduction

Actively growing cells have been shown to be about 17 to 26 percent protein by weight [1], with the implication that a similar fraction of a cell’s volume is occupied by protein. Additional volume is occupied by nucleic acids, ribosomes, and complex sugars. Together, these result in a very crowded intracellular environment that may be more physically similar to a protein crystal or gel than the dilute laboratory solutions that are typically used for in vitro experiments [1].

This macromolecular crowding affects intracellular dynamics in several ways (see reviews [2, 3, 4, 5, 6]). It slows diffusion by about a factor of 5 to 10 [7, 8], it increases association reaction rate constants by an order of magnitude or more [9], it favors folded protein conformations over unfolded ones [5], and it enhances the activity of chaperones [3, 10]. Most theoretical explorations of these effects have focused on thermodynamic considerations, including particularly the reduced translational and configurational entropies of molecules that are in crowded systems [11, 12]. This entropy reduction tends to decrease the entropic benefits of dissociated states and unfolded proteins, which then shifts equilibria toward associated states and folded proteins.

The effects of crowding can also be interpreted with a kinetic viewpoint. For bimolecular reactions, the reduction of available volume confines the reactants into less space, so they collide with each other more often than they would otherwise, which increases reaction rates. Vice versa, crowding also inhibits molecular motion, which decreases the rate at which reactants collide with each other and thus decreases reaction rates. Minton described these opposing effects of crowding on bimolecular reaction rates in 1990, predicting that volume reduction would be more important at low crowding densities and diffusion reduction at high crowding densities [13]. These predictions are qualitatively supported by recent experiments [9, 14, 15, 16, 17, 18, 19, 20] and modeling [21, 22, 23, 24]. The only quantitative theory of reaction rates in crowded environments has been reported so far is a model by Berezhkovskii and Szabo [25], although it stopped short of a complete description in terms of the system’s chemical composition.

This work presents two new quantitative models that account for the combined effects of volume reduction and diffusion inhibition on bimolecular reaction rates, as well as the effect of crowders on reducing reactants’ access to each other. This last effect is motivated by a simpler system, but one which is also important on its own merits, which is a semi-infinite space that is bounded on one side by an inert planar surface. Molecules near this surface are shown to react more slowly than those far from it. The “crowder proximity model” addresses reduced reactant access at relatively far distances from crowders, and the “reactant occlusion model” addresses reduced reactant access very close to crowders. This work also presents a completion to Berezhkovskii and Szabo’s theory, here called the “cavity model”. All three models agree comparably well with simulation data that were computed with the Smoldyn software, which has been thoroughly validated in prior work [26, 27, 28]. Each model also helps elucidate the physics in useful ways.

## 2. Theory

### 2.1. Reaction rates without crowders

This work builds on ideas introduced by Smoluchowski [29] and extended by Collins and Kimball [30]. For the generic irreversible reaction A + B → P, not including crowders, they showed that the rates of diffusion-influenced reactions can be computed using the radial distribution function of B molecules about A molecules, given here as *g*_*B*_(*r, t*). Doing so effectively defines a reference frame in which an A molecule is permanently at the origin and surrounded by B molecules that diffuse, in this reference frame, with the sum of the diffusion coefficients for the separate A and B molecules, *D*_*A*_ and *D*_*B*_; this sum is called the mutual diffusion coefficient and denoted *D* [31]. The A and B reactants contact each other when their centers are separated by the “contact radius”, *σ*_*AB*_, which equals the sum of the reactants’ physical radii, *σ*_*A*_ and *σ*_*B*_ (all molecules are assumed here to be spherical and rotationally isotropic, to diffuse ideally, and to only interact upon contact; see Figure 1A). For simplicity, it is sometimes helpful to assign the contact radius entirely to the A molecule, effectively making it a sphere of radius *σ*_*AB*_ and the B molecules into points (Figure 1B); this is valid whenever the A and B concentrations are sufficiently dilute that crowding interactions with other molecules of the same species can be ignored, which we assume here.

**Figure 1.**
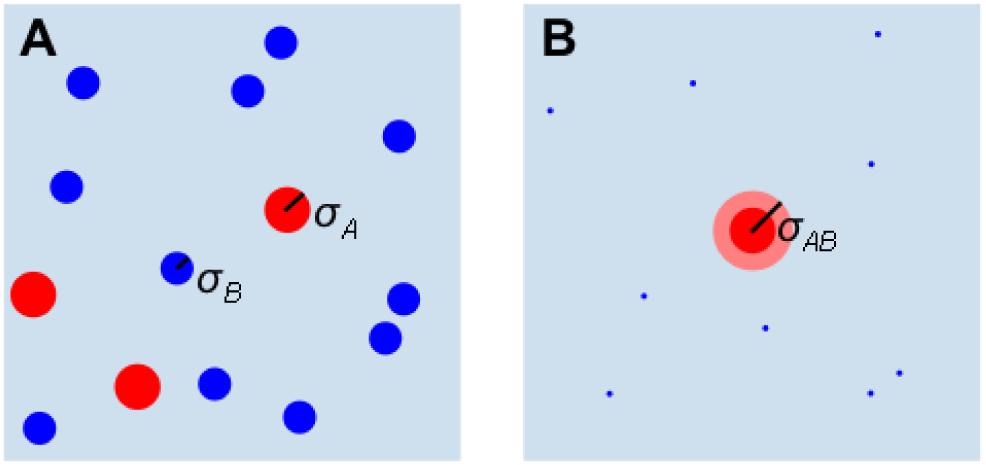
(A) A reactive system with A molecules in red and B molecules in blue, with respective radii *σ*_*A*_ and *σ*_*B*_. (B) A simplified version of the same system, now with an A molecule at the origin acting as a sink that has radius *σ*_*AB*_ and B molecules reduced to points.

The radial distribution function is normalized to approach 1 at large radii and evolves according to the radially symmetric diffusion equation [32],

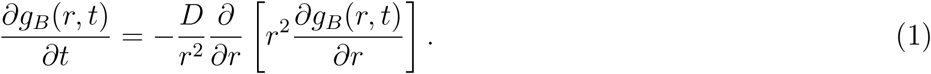

It has an inner Dirichlet boundary condition, *g*_*B*_(*σ*_*AB*_, *t*) = 0, if reaction occurs immediately upon contact as in the Smoluchowski model [29], or an inner Robin boundary condition,

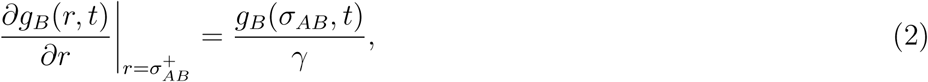

if contacting molecules have finite reactivity, as in the Collins and Kimball model [30]. Here, *γ* represents surface reflectivity, ranging from 0 for molecules that react upon contact, to infinity for molecules that never react. A finite reactivity can arise from a reaction activation energy or can represent orientational specificity for the reactants. It is often called the intrinsic reaction rate constant [31, 33], defined as 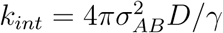.

The total reaction rate constant is the net influx of B molecules toward A molecules measured at the contact radius,

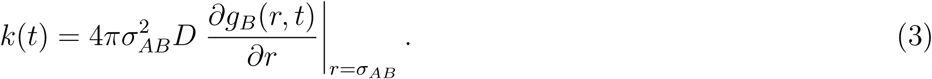

Solving eq. 1 for the steady-state radial distribution function, *g*_*B*_(*r*), and then using it to compute the reaction rate constant with eq. 3 leads to the well-known Collins and Kimball reaction rate equation [30, 34],

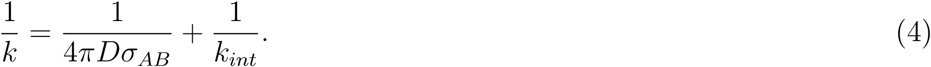

The first term represents the time that reactants take to initially find each other through diffusion and the second is the additional time that they then take to react. Reactions are said to be diffusion-limited if the first term dominates and activation-limited if the second term dominates. Alternatively, the degree to which a reaction is diffusion-limited can be quantified with the “diffusion-limited fraction”, *χ*, defined as

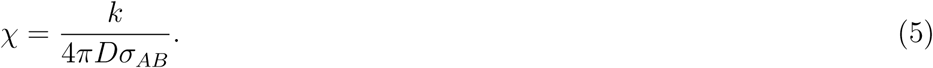

This value is 0 in the activation-limited extreme and 1 in the diffusion-limited extreme. It relates to *γ* as *χ* = *σ*_*AB*_/(*σ*_*AB*_ + *γ*).

### 2.2. Crowder volume exclusion and diffusion reduction

Now considering crowders, suppose they have number density *ρ*_*C*_, are spherical, are all the same size with radius *σ*_*C*_, and cannot overlap each other (Figure 2A). Minton predicted that crowding accelerates reactions by reducing available volume, and also slows reactions by slowing diffusion [13]; these effects are considered in turn.

**Figure 2.**
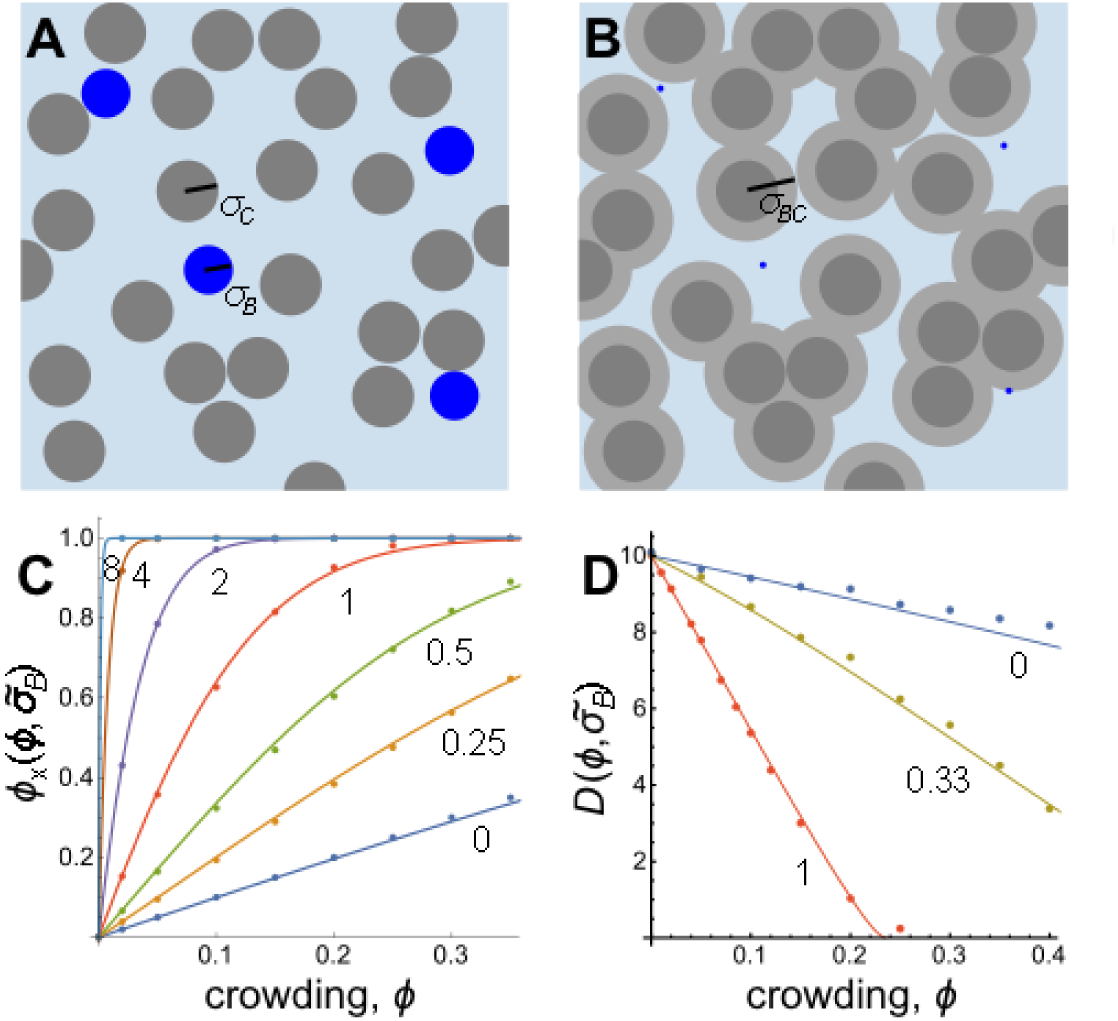
(A) Diagram of a crowded system, shown with B molecules and gray crowders with respective radii *σ*_*B*_ and *σ*_*C*_ ; crowders occupy volume fraction *ϕ*. (B) An equivalent system showing the B molecule centers and the total volume excluded by the crowders, now with effective radii *σ*_*BC*_ = *σ*_*B*_ + *σ*_*C*_ ; they exclude volume fraction 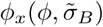. (C) Relationship between occupied and excluded volume fractions for the 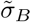 values shown. (D) Diffusion coefficients for B molecules with the 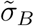 values shown, with *D*_0_ = 10 µm^2^/s.

The fraction of system volume occupied by crowders is the product of the volume for one crowder and the crowder density,

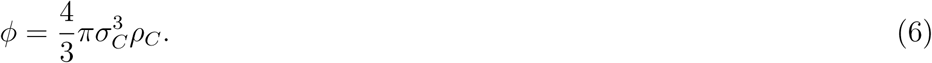

However, not all of the remaining volume is accessible to the centers of the reactant molecules because the reactants have finite radii that keep their centers away from the crowder edges [35] (Figure 2B). Focusing on the B molecules (the A molecules are identical, but we choose B molecules for concreteness), define the fraction of the total system volume that is excluded to the B molecule centers as 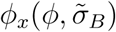. The tilde symbol implies a reduced radius in which the radius is divided by the crowder radius, so 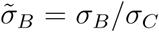. Clearly, 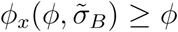 with equality only for molecules that are point particles. Figure 2C shows the relationship between 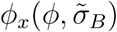 and *ϕ* with different curves representing different 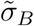 ratios. The points in the figure represent simulation data generated with the SmolCrowd software [27], which creates systems of randomly positioned spherical crowders that have non-overlapping core regions and overlapping outer regions, and then quantifies the excluded volume (see Methods). The curves in the figure were fit to the simulation data using an empirical function form, giving the relationship

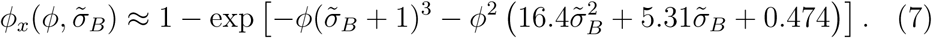

This fit is within 4% of all simulated values, which extend over all biologically important crowding fractions and all that are considered in this work, and can be shown to approach exactness in the limits of small *ϕ* and/or large 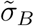.

The law of mass action states that reaction rates are directly proportional to each of the reactant concentrations [36], obeying the equation

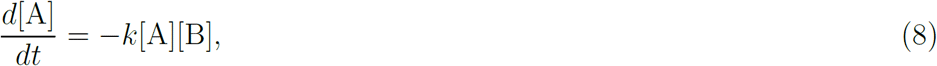

where brackets indicate concentrations. Assume that this applies within the portion of the system to which the A and B molecules are confined by the crowders; take this for now as the fraction 1 −*ϕ*_*x*_(*ϕ*) of the system volume. The observed reaction rate is always measured with respect to the total system volume rather than the accessible volume, so the concentrations in eq. 8 need to be corrected to account for the different volumes. Doing so introduces a factor of [1 − *ϕ*_*x*_(*ϕ*)]^−1^ on the right hand side of the equation, implying that the crowders increase the observed reaction rate constant by this factor; this is the reaction rate increase for Minton’s theory [13].

A similar analysis applies if the A and B molecules have different radii. Assume that the B molecules have the smaller radii, so that their accessible region of the system volume, *V* (1−*ϕ*_*B*_), is a superset of that which is accessible to the A molecules, *V* (1−*ϕ*_*A*_), where *V* is the total system volume and *ϕ*_*A*_ and *ϕ*_*B*_ are shortened versions of 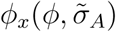 and 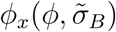. Reactions only occur in the portion of the volume that is accessible to both the A and B molecules, which is the region accessible to the A molecules, so eq. 8 becomes

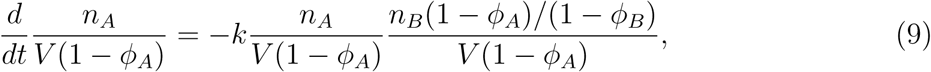

where *n*_*A*_ and *n*_*B*_ are numbers of A and B molecules, and the numerator of the last factor gives the number of B molecules that are in the volume that is accessible to A molecules. This simplifies to become the same as eq. 8, now considering concentrations relative to the entire system volume, but with the reaction rate constant increased by (1 −*ϕ*_*B*_)^−1^. This means that the effective volume for the reaction is the available volume for the less confined reactant. For convenience, we define *ϕ*_*x*_(*ϕ*) as the excluded volume for the less confined of either the A or B molecules.

The crowders reduce diffusion coefficients, shown in Figure 2D, because they physically block many diffusive trajectories. Points in this figure were computed by simulating diffusion of tracers, which had the same radii as B molecules but did not participate in reactions, among immobile spherical crowders using the Smoldyn software. Diffusion coefficients were then calculated from final mean square displacement values (see Methods). The lines in the figure, which agree well with the simulation data, were fit to a comparable data set in prior work [37]. They represent the function

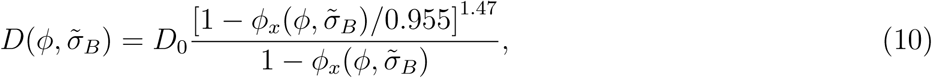

where *D*_0_ is the diffusion coefficient in the absence of crowders. This equation applies for 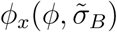 from 0 to 0.955; the upper bound is the excluded volume fraction where diffusion becomes completely stopped, which is called the percolation threshold. Although not explored in this work, note that diffusion coefficients are substantially larger if the crowders are mobile [38].

Combining the effects for the reduced available volume and slowed diffusion coefficients modifies the Collins and Kimball reaction rate equation from eq. 4 to

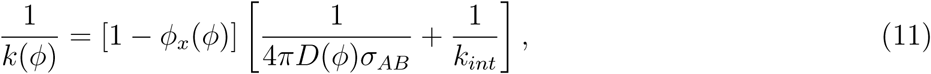

where *D*(*ϕ*) is the mutual diffusion coefficient in the crowded environment, which is equal to 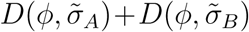. Figure 3 compares this prediction with simulation data for diffusion-limited and nearly activation-limited reactions in crowded systems. Predictions and data agree in the limits of no crowding at all (*ϕ* = 0) and so much crowding that diffusion completely stops (the percolation threshold, *ϕ* ≈ 0.25). However, they disagree strongly elsewhere, showing that the two effects included in this theory are inadequate to quantitatively explain how crowding affects reaction rates.

**Figure 3.**
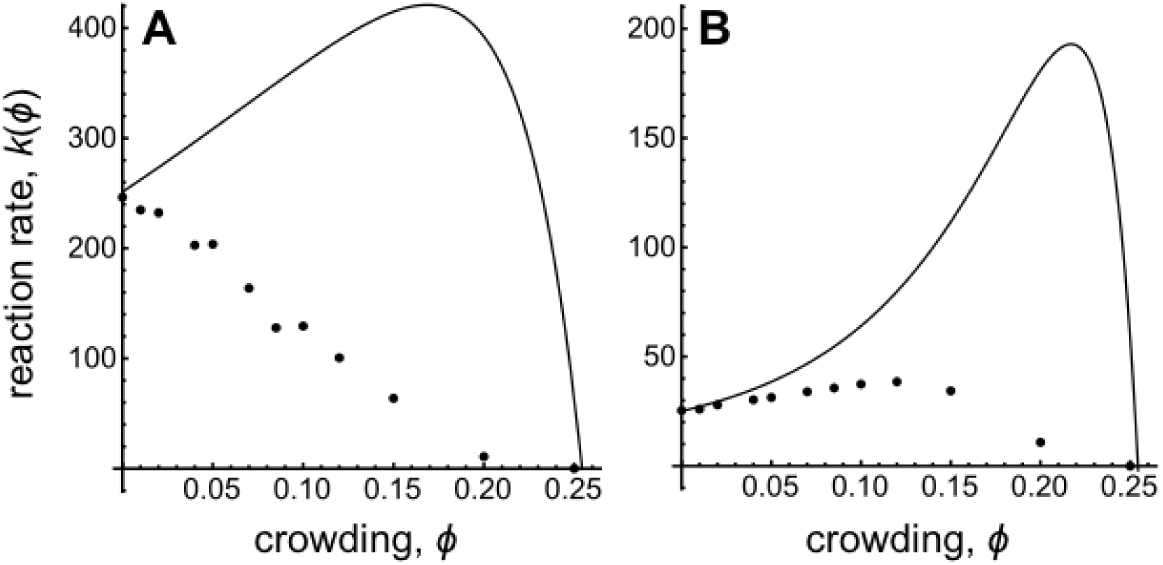
Effective reaction rate constants for (A) diffusion-limited and (B) nearly activation-limited reactions. Points are from Smoldyn simulations and lines from eq. 11. Parameters: *σ*_*A*_ = *σ*_*B*_ = *σ*_*C*_ = 0.5 nm and *D*_*A*_ = *D*_*B*_ = 10 µm^2^/s for both panels, and *χ* = 1 in panel A and 0.1 in panel B.

### 2.3. Reaction inhibition by a planar surface

A third way that crowding affects reaction rates is by reducing access to reactants. To investigate this effect in isolation, we consider the reaction A + B → P in semi-infinite uncrowded space, bounded on one side by an impermeable planar surface that is located at *x* = 0. A single A molecule sink is fixed at distance *x*_*A*_ from this surface. B molecules diffuse with diffusion coefficient *D*, and get absorbed by the sink (Figure 4A). We wish to solve for the steady-state reaction rate constant while accounting for the surface’s influence. This problem does not address crowding directly but is a useful intermediate step, as shown in subsequent sections.

**Figure 4.**
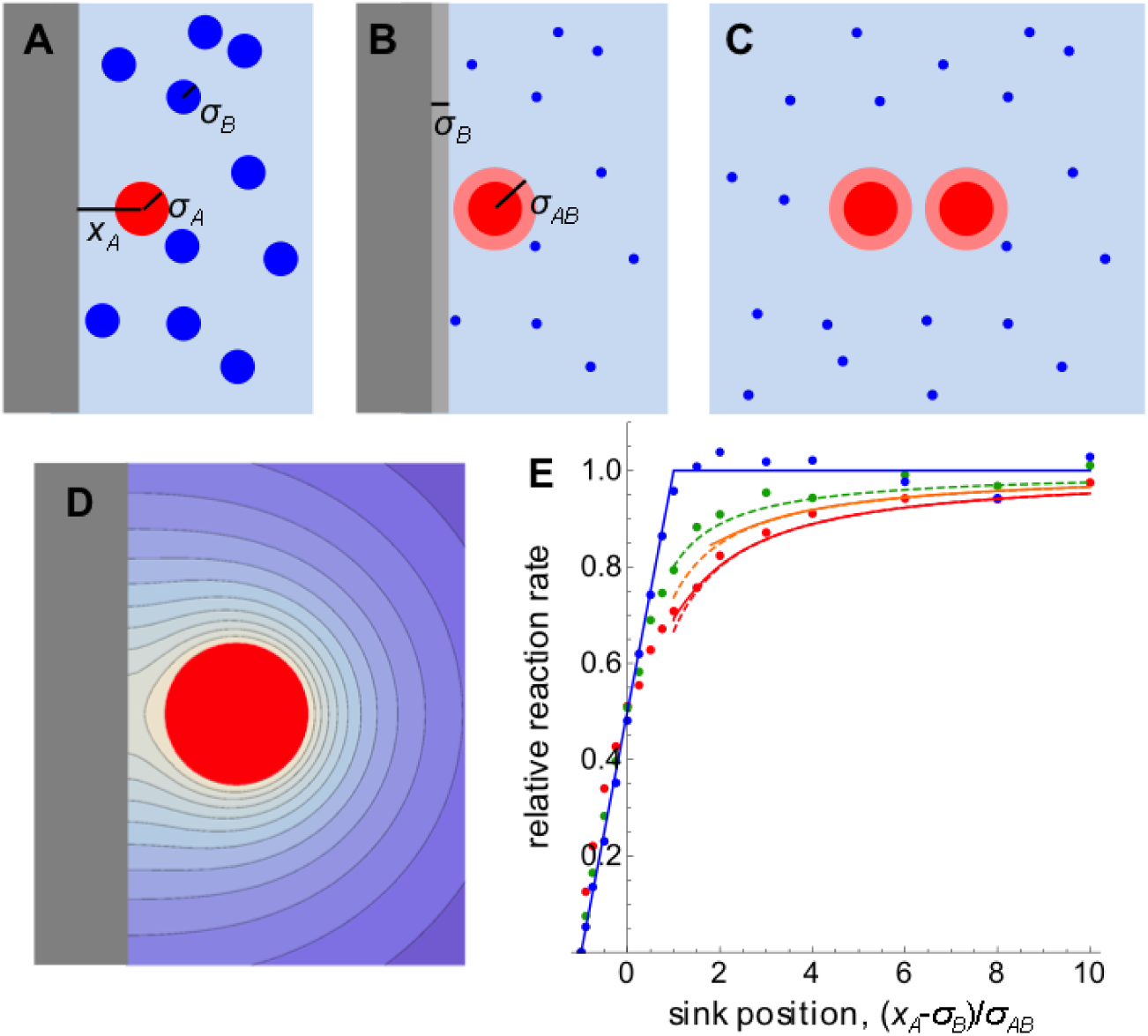
(A) A sink, in red, near an impermeable surface, in gray. B molecules, in blue, get absorbed by the sink. (B) A simplified version of the same system, with the B molecule radii effectively added to the surface and sink. (C) A further simplification using the method of images. (D) Contour plot of steady-state B molecule concentration about a sink. Note that contours are perpendicular to the surface where they touch, as required by the reflecting boundary condition. (E) Relative reaction rate as a function of distance from the surface. Red represents diffusion-limited (*χ* = 0.95 for the simulation data), orange is for *χ* = 0.7, green is for *χ* = 0.5, and blue is for activation-limited (*χ* = 0.09 for the simulation data). Points are from Smoldyn simulations, solid lines from theory, and dashed lines from eq. 14.

As usual, it’s convenient to focus on the centers of the B molecules rather than their edges, which effectively moves the surface location to *σ*_*B*_ and increases the A molecule radius to *σ*_*AB*_ (Figure 4B). Define *C*_*B*_(**x**) as the steady-state mean B molecule concentration at position **x**. It approaches the overall B molecule concentration at large distances from the A molecule but is smaller at closer distances due to absorption. The system has a Neumann boundary condition at the effective planar surface,

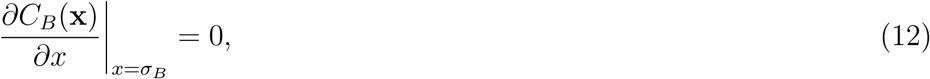

to represent the fact that there cannot be concentration gradients perpendicular to impermeable surfaces. As before, the sink has a Robin boundary condition at the contact radius, *σ*_*AB*_, to represent reactions (eq. 2). The Neumann boundary condition at the planar surface is easiest to address with the method of images [39]. Here, this means that a mirror image of the real system, including the A molecule sink, is introduced on the unphysical side of the surface. By symmetry, this modified system automatically obeys the Neumann boundary condition at *x* = *σ*_*B*_, allowing the presence of the surface to be neglected (Figure 4C).

The steady-state concentrations (Figure 4D) and reaction rates have been solved exactly for this symmetric system, both for the case where reactants react immediately upon contact [40] as in the Smoluchowski model, and where they have finite reactivity [41] as in the Collins and Kimball model. Both solutions require that the sink does not touch the planar surface (i.e. *x*_*A*_ *> σ*_*AB*_ + *σ*_*B*_), which I call the unoccluded case. The former reaction rate is [40]

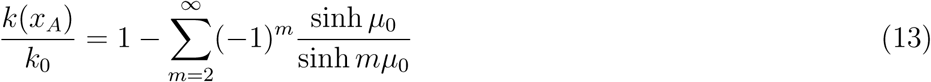

where *k*_0_ is the reaction rate constant without the surface and the *µ*_0_ parameter is determined from cosh *µ*_0_ = (*x*_*A*_ −*σ*_*B*_)/*σ*_*AB*_. The latter reaction rate, for the Collins and Kimball model, is not repeated here because it is complicated, but is described in ref. [41]. Figure 4E shows these analytic solutions with red and orange solid lines. Both solutions are closely approximated by

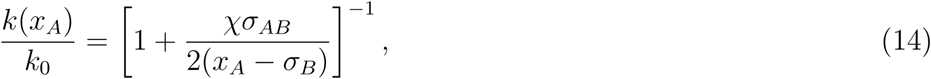

shown in Figure 4E with dashed lines. This equation, which I derived using a particle emission model as described below, extends a result for the Dirichlet boundary condition that is given in ref. [40] and, again, only applies for the unoccluded case. Both the exact and approximate solutions agree well with simulation results, shown in the figure with points.

The occluded case occurs when the sink overlaps the planar surface, meaning that the A molecule edge is within a B molecule diameter of the surface. This is harder to address but a few solutions can be found. (1) When the center of the A molecule is exactly at the effective surface, meaning *x*_*A*_ = *σ*_*B*_, the actual A molecule sink exactly overlaps its image to make a single sphere, so the reaction rate per sink is half of that for a single isolated sink. This means that *k*(*σ*_*B*_)/*k*_0_ = 1/2. (2) The sink is inaccessible when it is fully behind the surface, so it must have zero reaction rate. This implies that *k*(*x*_*A*_) = 0 for *x*_*A*_ ≤ −*σ*_*A*_ or, equivalently, for *x*_*A*_ − *σ*_*B*_ ≤ −*σ*_*AB*_. (3) In the activation-limited extreme, *C*_*B*_(**x**) is the same everywhere, so absorption at the sink depends only on the amount of exposed sink surface area. This means that *k*(*x*_*A*_)/*k*_0_ = 1 in the unoccluded case and *k*(*x*_*A*_)/*k*_0_ = (*x*_*A*_ + *σ*_*A*_)/2*σ*_*AB*_ in the occluded case (this is easily derived from the surface area of a “spherical cap”, which is 2*πrh* where *r* is the sphere radius and *h* is the cap height). Figure 4D shows this last solution with a blue line.

This analysis shows that the mere presence of an impermeable surface can substantially decrease the reaction rate constant for nearby molecules, even if they are several radii away from the surface. This reduced reactivity arises because the surface reduces access to the side of a molecule that faces the surface. This is apparent in Figure 4D, where the steady-state B molecule concentration is low between the A molecule and the surface and, further, has a very shallow gradient, which then implies a low reaction rate (see eq. 3). As another view of the same phenomenon, each A molecule is effectively competing with its mirror image for B molecules, thus causing it to absorb fewer B molecules than it would when far from a surface. This decreased reactant access is especially acute in the occluded case, where part of one reactant is completely inaccessible.

The finding that reaction rates are slower near surfaces raises the intriguing possibility that cells might control some reaction rates by regulating the precise locations of reactive sites that are near membranes. For example, cells could extend or retract specific membane-bound receptors by small amounts to modulate their rates of ligand binding. Also, cells could reorient protein binding sites toward or away from membranes to change their reactivities, using the different gradients at the sink surface that are shown in Figure 4D. I am not aware of examples where there is evidence of these but the necessary protein conformational changes are well within the realm of typical biochemical behaviors.

At least in principle, surfaces can also accelerate reaction rates if molecules can adsorb and then diffuse along the surface, due to reduction of dimensionality effects [42, 43]. For example, if the B molecules in Figure 4 could reversibly adsorb to the surface, and then diffused much faster along the surface than they do in solution, then this would draw high B molecule concentrations in close to the sink, leading to a fast reaction rate. In practice, this is likely to be a weak effect because adsorbed molecules typically diffuse substantially more slowly than free molecules [44, 28].

### 2.4. Crowder proximity model

Given that proximity to a planar surface slows reaction rates by reducing reactant access to each other, it makes sense that proximity to the multiple surfaces of crowded systems would slow reaction rates in the same way. This is an aspect of diffusion inhibition by crowders but is independent of the slower diffusion that was considered previously in eq. 11. Ideally, we would address this effect by computing the influx rate for a sink that is surrounded by randomly placed impermeable spheres.

However, this is impossible because eq. 13 does not generalize to spherical surfaces. Thus, we use an approximate method instead, in which we invert the problem to consider emission of B molecules out of a point source instead of absorption of B molecules into a sink. This inverted problem obeys the same diffusion equation and has the same Neumann boundary conditions at the crowder surfaces but has net outward flow instead of net inward flow, and the concentration approaches zero at large distances instead of a finite value. The only substantial change is that the Robin boundary condition at the contact radius (eq. 2) is ignored for now.

Consider an emitter at the origin that emits B molecules at unit rate, and a spherical crowder that has its center at distance *r*_0_ and has radius *σ*_*C*_ (Figure 5A). We focus on the centers of the B molecules, giving the crowder an effective radius of *σ*_*BC*_ = *σ*_*B*_ + *σ*_*C*_. Dassios and Sten showed that the Neumann boundary condition at the effective crowder surface can be addressed with the method of images [45], much as before. In this case, there is a “point image emitter” that is within the crowder and at distance 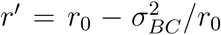 from the origin, which emits at rate *q*_*pt*._ = *σ*_*BC*_/*r*_0_, and also a “line image emitter” that extends from *r′* to *r*_0_ and emits at a continuous rate over its entire length with emission density *q*_*line*_ = −1/*σ*_*BC*_. The line emitter emits at a negative rate, which is mathematically sensible as stated, or can be considered physically as emission of anti-particles that anhiliate with normal particles upon collision. The important aspect of these image emitters is that adding them to the system creates the same B molecule concentration as occurs with the crowder at all positions outside of the crowder’s effective surface, but without requiring the crowder itself. In particular, the concentration gradient at the effective crowder surface is zero (Figure 5B), as required. Concentrations are different inside the crowder, but these are unphysical and so can be ignored.

**Figure 5.**
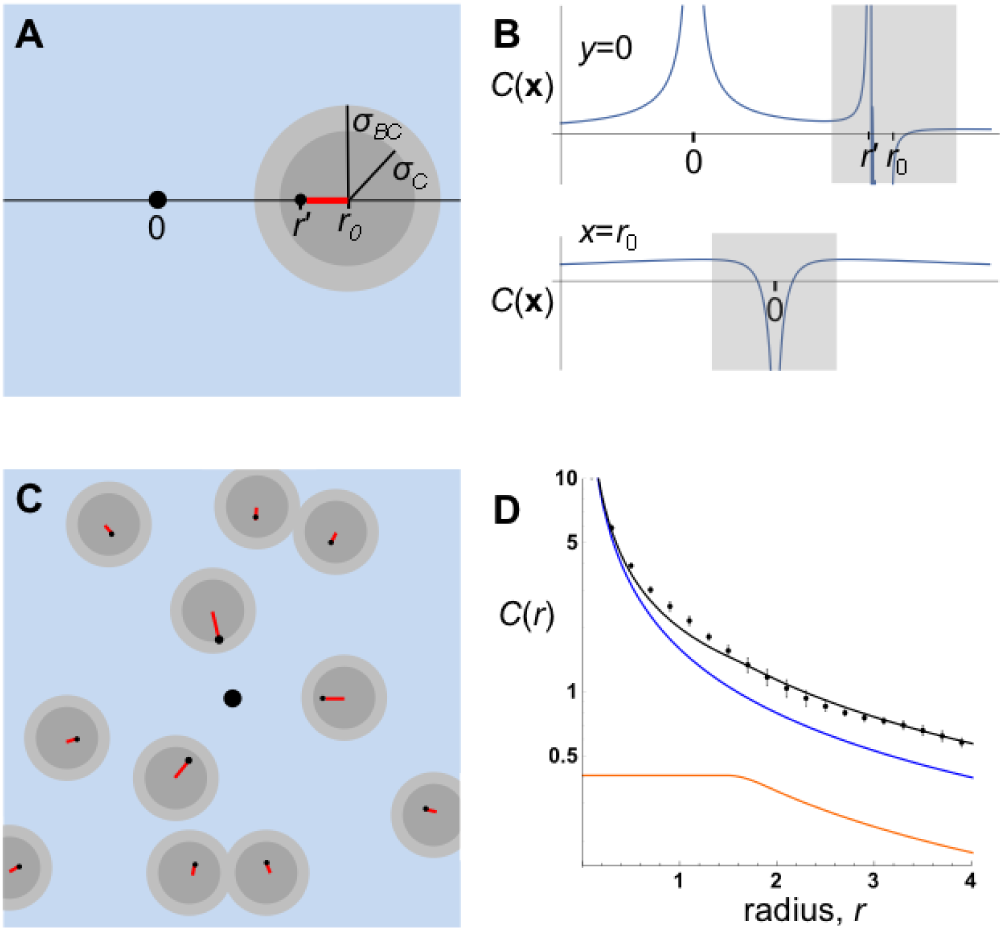
(A) An emitter at the origin, shown at *x* = 0 in black, and its image emitters within a spherical crowder, shown as a black dot at *x* = *r′* for the point emitter and as a red line for the line emitter. (B) The total emitted B molecule concentration measured along the *x*-axis at top and along *y* at *x* = *r*_0_ on the bottom. Dark bands represent the crowder, at the edges of which the Neumann boundary condition is observed; internal concentrations are unphysical. (C) An emitter at the origin surrounded by randomly positioned crowders, each of which has point and line emitters. (D) B molecule concentration as a function of radius for the system shown in panel C, for *ϕ* = 0.3. The blue line represents the concentration from just the central emitter, orange represents the concentration from the image emitters, and black represents the total concentration. Dots represent data from Smoldyn simulation using impermeable spheres rather than image emitters. Data were averaged over 10 random crowder configurations; error bars show 1 standard deviation.

We extend this image emitter solution to many crowders (Figure 5C) using a mean field approximation, replacing discrete point and line image emitters for individual crowders with a density of point and line image emitters for a density of crowders. In the process, we only consider images that arise from the B molecule emitter at the origin, while ignoring higher order images of images. Normalized to an overall crowder density of *ρ*_*C*_ = 1, the density of emission at radius *r′* from the origin that arises from point image emitters is

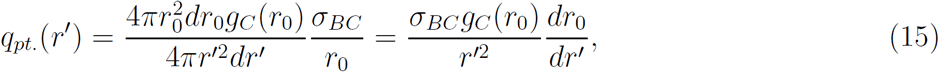

where *g*_*C*_ (*r*_0_) is the radial distribution function of crowders about the origin. In the first equality, the numerator of the first term gives the number of crowders that have their centers between *r*_0_ and *r*_0_+*dr*_0_, the second term is the emission rate from these crowders, and the denominator of the first term converts the result back to a density at radius *r′*. The *r*_0_ values in this emission density can be replaced using 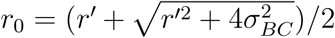, from the relationship between *r*_0_ and *r′* given above, but this only complicates the result. The density of emission at radius *r′* from line image emitters requires integration over all of the crowders that contribute to line image emission at this location, leading to

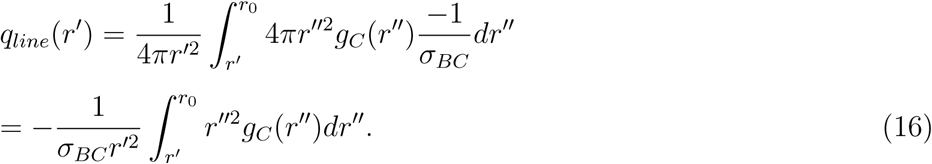

In the first equality, the integrand represents the line emission at *r′* by crowders centered between *r″* and *r″* + *dr″*, it is integrated over all crowders that contribute emission here, and the initial factor converts the result to a density at *r′*. The total density of emission at *r′*, still normalized to unit crowder density, is the sum of these contributions,

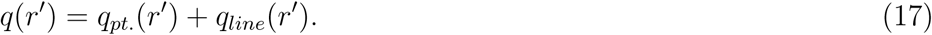

This image emitter density depends only on the crowder locations and effective sizes, so it can be evalulated from only knowledge of *g*_*C*_ (*r*_0_) and *σ*_*BC*_, meaning that details about A molecules, reactions, and all concentrations are unimportant here.

Next, we compute the steady-state B molecule concentration, from both the emitter at the origin and all image emitters. We do so with a Green’s function approach, using the fact that the steady-state density of emitted particles from a thin spherical shell of emitters that is located at radius *r*_*s*_ from the origin is [32]

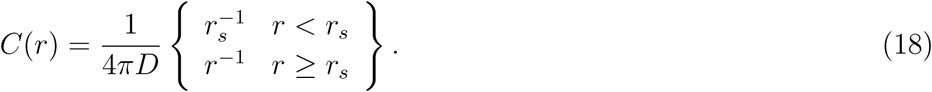

Integrating this kernel over the actual density of emitters, including the emitter at the origin and all of the image emitters (eq. 17), gives the average concentration of B molecules as

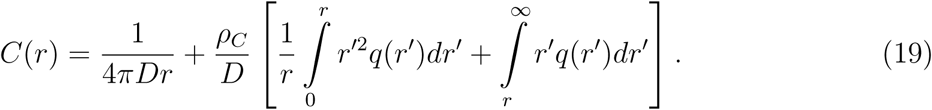

Figure 5D shows the emitted particle density for the case where *σ*_*BC*_ = 1 and *g*_*C*_ (*r*_0_) is a step function that is 0 for *r*_0_ *<* 2 and 1 for *r*_0_ *>* 2. The blue line represents the first term of eq. 19 (from the emitter at the origin), the orange line represents the second term (from the image emitters), and the black line represents their sum. Points in the figure show that simulation results, which were computed with randomly placed impermeable spheres rather than image emitters, agree well with the theory.

Some transformations convert eq. 19, which is for B molecule emission from a source at the origin, back to the desired radial distribution function for a reactive sink at the origin. The sign is reversed to change from emission to absorption, the concentration is rescaled so that it obeys the desired Robin boundary at the contact radius (which can now be obeyed due to the spherical symmetry introduced by the mean field approach), and the result is offset so that the radial distribution approaches 1 at large radii. Together these give

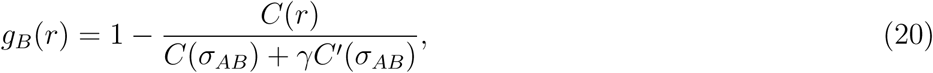

where 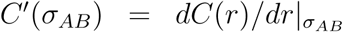. For example, if there are no crowders, then *C*(*r*) = 1/4*πDr*, which transforms to the radial distribution function 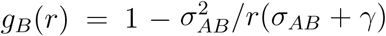, as it should be for the Collins and Kimball model [26]. Substituting this transformation into eq. 3 gives the steady-state reaction rate constant in terms of the concentration of emitted B molecules,

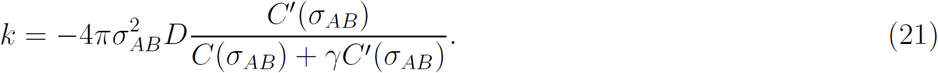

If there are no image emitters within radius *σ*_*AB*_, discussed below, then this reaction rate equation can be expanded using eq. 19 and then simplified to

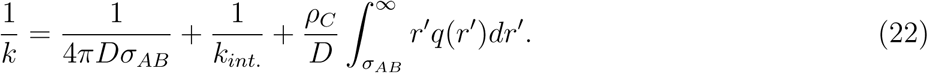

The first two terms are the same as those in the Collins and Kimball reaction rate equation, eq. 4, while the last term is new and accounts for inhibited reactant access from nearby crowder surfaces. Define *Q* as a scaled version of this new term,

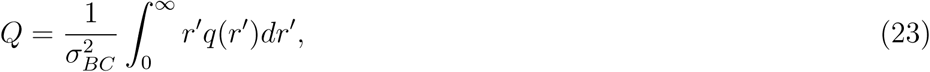

where the lower integration limit was reduced to 0 for simplicity but didn’t change the integral’s value because of the assumption of no image emitters within *σ*_*AB*_. This is a unitless number that quantifies the amount of reactant access decrease that arises from nearby crowders.

Eq. 22 is based on image emitters rather than actual crowders, so it does not account for either the excluded volumes of the crowders or their effects on diffusion coefficients. We include these effects in the same way that we did for eq. 11 to give a final reaction rate equation for irreversible reactions in crowded environments,

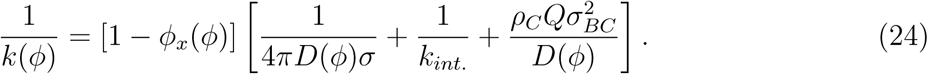

This is the complete equation for the reaction rate in what I call the “crowder proximity model”, now accounting for crowder volume exclusion, diffusion inhibition, and inhibited reactant access.

We still need to compute *Q*, which comes from the crowder radial distribution function, *g*_*C*_ (*r*_0_). Ignoring the structured layers that arise in densely packed hard spheres [46], the obvious first-order approximation is that crowders should be well mixed but cannot overlap A molecules. In other words, *g*_*C*_ (*r*_0_) should be a step function that is 0 for *r*_0_ *< σ*_*AC*_ (where *σ*_*AC*_ = *σ*_*A*_ + *σ*_*C*_) to account for the excluded volumes of the A molecules and crowders, and 1 for larger *r*_0_ values,

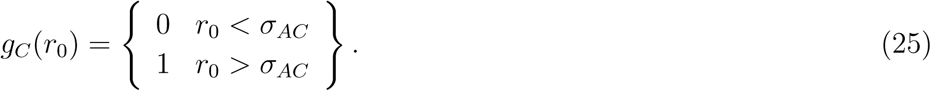

However, this doesn’t work in the theory developed here because a crowder at *σ*_*AC*_ would have a point image emitter at 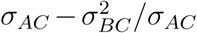, which may be inside of *σ*_*AB*_; this violates an assumption made when deriving eq. 22. Physically, the problem is that the theory only works for the unoccluded case in which the sink, which has radius *σ*_*AB*_, does not overlap the effective surface of the crowder, which has radius *σ*_*BC*_. In other words, each A molecule must be far enough from each crowder that a B molecule can fit between them. To address this problem, we move the step in the crowder radial distribution function, eq. 25, outward to *σ*_*ABBC*_ = *σ*_*A*_ + 2*σ*_*B*_ + *σ*_*C*_. This now satisfies the assumption of the unoccluded case, but we recognize that the result will underestimate the amount of reactant access inhibition. With this change, *Q* evaluates to

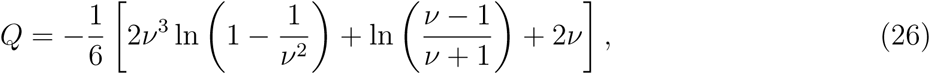

where *ν* = *σ*_*ABBC*_/*σ*_*BC*_. For example, if *σ*_*A*_ = *σ*_*C*_, then *ν* = 2 for all *σ*_*B*_ values, and *Q* = 0.283.

Red lines in Figure 6 compare the theoretical reaction rate constant from eq. 24 with simulation data, shown with points. Agreement is reasonably good for low to medium crowding densities over a wide range of system parameters, including different relative sizes of *σ*_*A*_, *σ*_*B*_, and *σ*_*C*_, and different amounts of diffusion influence on reactions. However, as expected, the theory underestimates reactant access inhibition at high crowding densities, yielding reaction rate predictions that are too fast.

**Figure 6.**
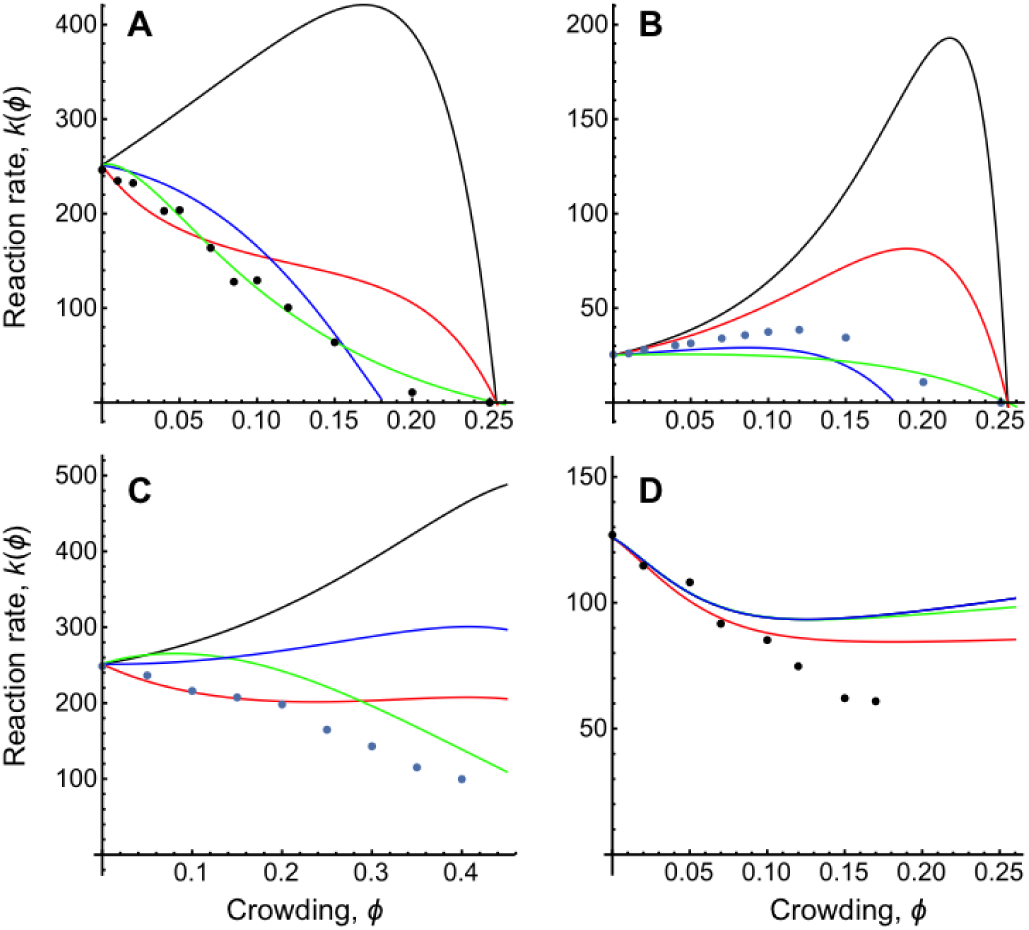
Reaction rates as a function of crowding for different parameter values. Points represent simulation data, black lines represent a model that ignores crowding effects on reactant access (eq. 11), red lines represent the crowder proximity model (eq. 24), blue lines represent the reactant occlusion model (eq. 31), and light green lines represent the cavity model (eq. 36). (A) Diffusion-limited reactions with all species the same size: *χ* = 1, *σ*_*A*_ = *σ*_*B*_ = *σ*_*C*_ = 0.5 nm. (B) Nearly activation-limited reactions with all species the same size: *χ* = 0.1, *σ*_*A*_ = *σ*_*B*_ = *σ*_*C*_ = 0.5 nm. (C) Diffusion-limited reactions with larger crowders: *χ* = 1, *σ*_*A*_ = *σ*_*B*_ = 0.5 nm, *σ*_*C*_ = 1.5 nm. (D) Diffusion-influenced reactions with point-like B molecules: *χ* = 0.5, *σ*_*A*_ = 1 nm, *σ*_*B*_ = 0 nm, *σ*_*C*_ = 0.5 nm. In all cases, A and B diffusion coefficients were 10 µm^2^/s and their interaction radius, *σ*_*AB*_, was 1 nm.

### 2.5. Reactant occlusion model

There is no simple way to account for the effects of reactant occlusion (Figure 7A) for general reactions. However, an approximation is possible if the reaction is activation-limited because the rate is then directly proportional to the exposed effective surface area of the reactants, as shown above in section 2.3 (see the blue line in Figure 4E). We follow this approach here. As usual, we transfer the B molecule radii to the sink and crowders, giving them radii *σ*_*AB*_ and *σ*_*BC*_, respectively (Figure 7B).

**Figure 7.**
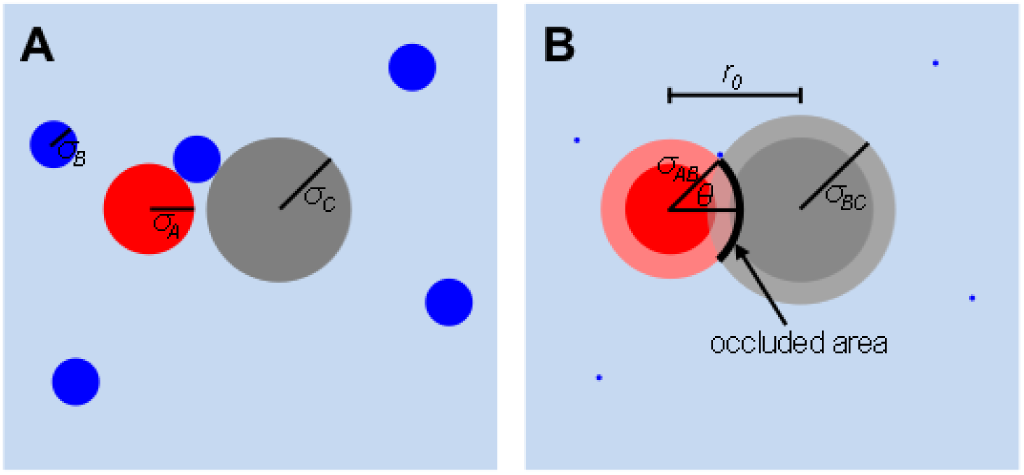
Reactant occlusion model. (A) An A molecule (red) that is close to a crowder (gray), with nearby B molecules (blue). The B molecule touching the A molecule and crowder cannot get further in between them. (B) The same molecules, but with B molecule radii transferred to the A molecule and crowder. The black “occluded area” is inaccessible to B molecule centers.

When a sink and crowder overlap, the occluded portion of the sink is a spherical cap. The area of a spherical cap is *A*_*cap*_ = 2*πr*^2^(1 − cos *θ*), where *r* is the radius and *θ* is the polar angle. In this case, *r* = *σ*_*AB*_ and *θ* can be found from the law of cosines, which together give the cap area as

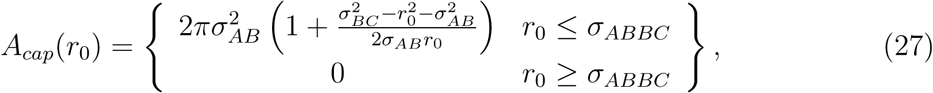

where *r*_0_ is the distance between the sink and crowder centers.

The average fraction of a sink’s surface area that is occluded, *A*_*occ*._/*A*_*total*_, can be estimated by adding the cap areas for a mean field of crowders, and then dividing by the total sink surface area,

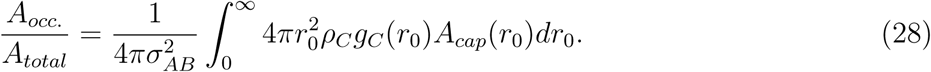

Substituting in eq. 25 for *g*_*C*_ (*r*_0_), eq. 27 for *A*_*cap*_(*r*_0_), integrating, and then simplifying yields

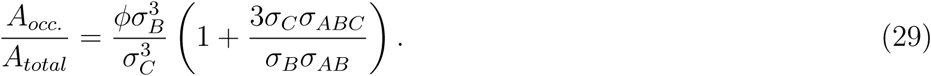

The mean field approximation is valid for low crowding densities but becomes inaccurate at high densities for two reasons. First, it ignores the restriction that crowders cannot overlap each other, with the result that it includes crowders at impossible locations. Second, reactant surface regions that are occluded by multiple crowders at once are included separately in the integral, despite the fact that an occluded region of the surface cannot be occluded again. These errors can be seen in the fact that eq. 29 is directly proportional to *ϕ*, causing it to predict that sufficient crowding can cause more than 100% of a sink to become occluded, which is obviously impossible. These limitations cause eq. 29 to overestimate the occluded surface area fraction.

For activation-limited reactions, the reaction rate simply scales with the accessible reactant surface area, making it equal to

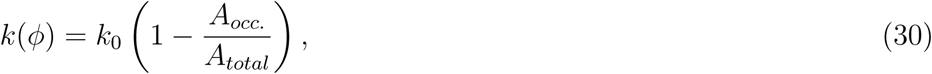

where *k*_0_ is the reaction rate without crowders. Combining eqs. 30, 11, and 29 yields the reaction rate predition for the “reactant occlusion model”,

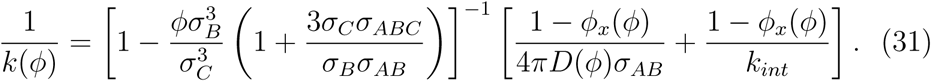

Figure 6 shows these predictions with blue lines. As expected, they are reasonably accurate for activation-limited reactions (Figure 6B) with low crowding and then underestimate reaction rates with high crowding. Also as expected, this model predicts that crowders don’t reduce access to reactants at all when the B molecules are point particles (Figure 6D); this overestimates reaction rates because it ignores the reduced access that arises in the unoccluded case. The model is surprisingly accurate for other diffusion-limited reactions (Figures 6A and 6C), where it was not expected to apply at all.

### 2.6. Cavity model

The previously published model for reaction rates in crowded systems, by Berezhkovskii and Szabo, considers the region between A molecule sinks and nearby crowders as a cavity [25] (Figure 8). Using the terminology and notation introduced above, the mutual diffusion coefficient is *D* within the cavity and *D*(*ϕ*) outside of it. The cavity also acts as a potential energy well that creates an effective attractive force between A and B reactants. The authors solved the diffusion equation for this model to give the reaction rate as

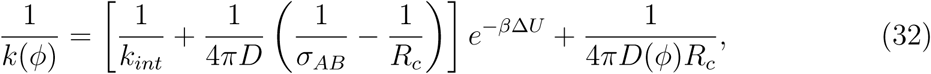

where Δ*U* is the energy well depth, *β* is 1/(*k*_*B*_*T*) in which *k*_*B*_ is Boltzmann’s constant and *T* is the temperature, and *R*_*c*_ is the cavity radius. They left Δ*U* and *R*_*c*_ as unsolved parameters. The authors also presented a more general model that accounts for molecule adsorption and diffusion on crowders, which we do not consider here.

**Figure 8.**
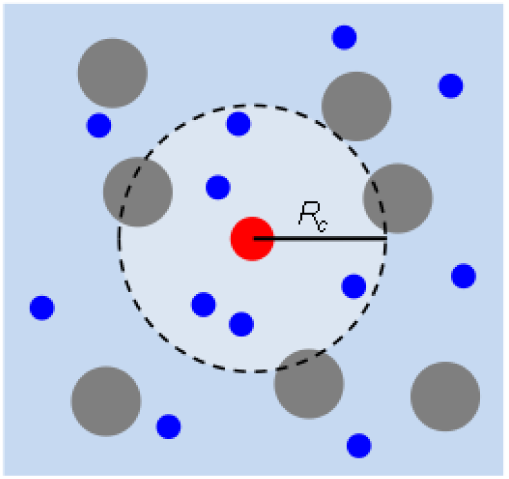
Cavity model. A cavity is defined about an A molecule (red) that extends out to the nearest crowders (gray). It acts as a potential energy well for B molecules (blue), which also diffuse faster within the cavity.

There is no actual energy difference between regions inside and outside of the cavity under the assumptions used here and in their work, but it is reasonable to consider a Helmholtz free energy difference,

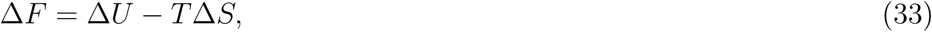

where Δ*S* is the entropy difference. Thus, we replace Δ*U* in eq. 32 with Δ*F*. Consider equal total volumes of crowded and uncrowded spaces, both given as *V*. A molecule that moves from the uncrowded space to the crowded space moves from an accessible volume of *V* to an accessible volume of *V* [1 − *ϕ*_*x*_(*ϕ*)]. If intermolecular forces can be ignored, as in an ideal gas, and the system is at constant temperature, then this transition corresponds to an entropy change of

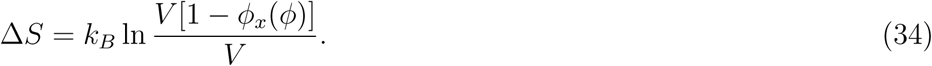

Under the assumption that the energy difference is 0, eqs. 33 and 34 combine to give

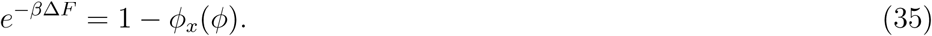

This reaction acceleration factor agrees with the result derived above from kinetic arguments.

Substituting this reaction acceleration factor into eq. 32 and rearranging leads to our “cavity model” prediction,

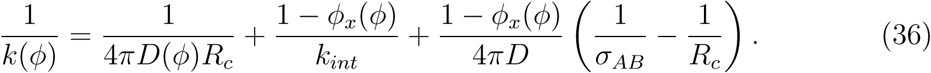

This equation is structurally similar to the crowder proximity model result, eq. 24, but has some important differences with regard to which terms include reaction acceleration factors and which use *D* versus *D*(*ϕ*). Here, the first term represents the time for a B molecule to diffuse to the edge of the cavity, the third term presumably represents the time to go from the cavity edge to the A molecule, and the second term represents the time to react after the initial collision.

We define the cavity radius, *R*_*c*_, as the mean radius around an A molecule within which there is one crowder or, equivalently, multiple parts of crowders that add to one. Using this definition and the crowder radial distribution function from eq. 25,

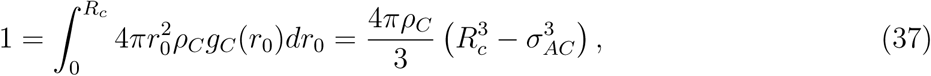

which rearranges to

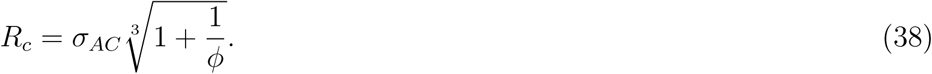

This radius expands as *ϕ*^−1/3^ for low crowding fractions and is on the order of *σ*_*AC*_ for high crowding fractions, both as expected [25].

Figure 6 shows this cavity model prediction with light green lines. Overall, it appears to be somewhat better than the other theories, but not substantially. Minor changes in the cavity radius definition, such as omitting the first term under the radical in eq. 38, did not change results appreciably.

## 3. Discussion

Minton described two opposing effects of macromolecular crowding on bimolecular reaction rates, which were that volume exclusion should accelerate reactions and diffusion inhibition should slow them. This work addresses the topic quantitatively and finds that a simple interpretation of these effects, eq. 11, disagrees with simulation results, substantially overestimating reaction rate constants. To investigate the possibility that this error arises from neglecting the effect of crowders on nearby reactants, this work considers reaction rates for molecules that are fixed near planar surfaces. Surface proximity is found to substantially slow reactions, given in eqs. 13 and 14, even when reactants are several radii away from the surface. It slows reactions even more if the surface is close enough to block access to part of a reactant, here called the occluded case. Applying this surface proximity effect to macromolecular crowding using a mean field approximation yields a more accurate reaction rate equation, eq. 24. This “crowder proximity model” agrees reasonably well with simulations for low crowding densities but still overestimates reaction rates at high densities because it only addresses the unoccluded case. A different “reactant occusion model”, eq. 31, takes the opposite approach by only considering reactant occlusion by crowders. This model also agrees reasonably with simulations at low densities but diverges at high densities, in part because it overcounts reactant occlusion. Finally, this work completes a “cavity model” that was introduced previously [25], yielding eq. 36, finding that it performs comparably with the other models.

Even if the crowder proximity model addressed the occluded case correctly, other approximations would likely cause errors, albeit smaller ones. (1) The mean field theory given in eq. 17 only includes images of the central emitter, while ignoring higher order images of images. The first order images in each crowder have positive emission toward the origin and negative emission away from the origin, so their images would add to the existing surface effect to make it stronger. This means that accounting for these effects would reduce predicted reaction rates. However, the total emission from each crowder adds to zero, making it unlikely that these higher order images would have a large effect. Further, simulation and theory agreed well for a central emitter (Figure 5D), again suggesting that this effect is minimal. (2) The mean field theory treats every crowder as an isolated sphere of the same size, ignoring the fact that the effective radii of multiple crowders can overlap each other to create larger obstacles (Figure 2B). These larger obstacles have less total surface area than the individual crowders, but are nevertheless more effective at reducing access to reactants because molecules can’t go between them anymore but instead have to diffuse a longer distance to get around them (note the 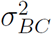 dependence in eq. 24). Accounting for this effect would also reduce predicted reaction rates. (3) The crowder radial distribution function, eq. 25, ignores reaction inhibition that is caused by crowders. This reaction inhibition decreases reaction rates close to the crowders, enabling reactants that are close to crowders to survive longer. As a result, crowders are statistically closer to reactants than assumed here. Again, accounting for this effect would reduce predicted reaction rates.

The reactant occlusion model has more serious flaws. It ignores the restriction that crowders can’t overlap each other, it counts occlusion by multiple crowders separately despite the fact that any portion of the reactant effective surface can only be occluded once, and it ignores reaction inhibition for the unoccluded case. If the first two issues could be fixed, then it would presumably be an accurate model for activation-limited reactions as, in fact, a large fraction of biochemical reactions are [31].

The more abstract nature of the cavity model makes it harder to analyze or improve upon, but it is nevertheless a promising theory for future work. Further, its simplicity makes it attractive for generalizing to a broader range of interactions, such as the original authors’ inclusion of reactant adsorption to crowders [25].

It would be nice to compare these theoretical predictions with experiment, but existing experimental data are unfortunately inadequate due to lack of breadth of parameters investigated, insufficiently characterized reaction dynamics, and/or presence of artifacts from non-specific binding or other contributions [12]. Instead, simulations are the only method currently available for producing data that can test these theories quantitatively. The Smoldyn software [26, 27, 47] is particularly well suited for these types of simulations because it is fast and accurate, has the necessary features (such as generating randomly distributed non-overlapping crowders and computing radial distribution functions), and does not impose a lattice on the system, which can introduce substantial artifacts [48]. Also, all of its algorithms approach exactness as simulation time steps are reduced toward zero, enabling one to make results as accurate as desired (here, time steps were always reduced until results stopped changing in order to achieve nearly exact results; see Methods). Green’s Function Reaction Dynamics (GFRD) [49, 50, 51, 52] methods would be an alternate approach, and have the benefit of being actually exact, but are impractical because they run several orders of magnitude slower than Smoldyn for systems with this level of complexity [24, 53]. Other options include the ReaDDy [54, 55] and SpringSaLaD [56] packages, both of which offer good accuracy but, again, tests showed that they run several orders of magnitude slower [53].

It would be straightforward to extend the first two models developed here to account for crowders with a distribution of sizes rather than the single size assumed here. In the crowder proximity model, each crowder in the distribution could still be represented with point and line image emitters, and the effects of those emitters would still add linearly (this approach also works for non-spherical crowders but would become much more complicated). At the end, the only change would be that the single crowder number density value, *ρ*_*C*_, would be replaced by a separate density for each crowder size or with a density function, *ρ*_*C*_ (*σ*_*C*_). Adding or integrating over these would then yield the surface effect term for eq. 24. Likewise, the reactant occlusion model could be extended by generalizing the integral in eq. 28 to work with multiple crowder sizes. The cavity model already applies to mixed crowder sizes (and non-spherical crowders).

On the other hand, it may be more difficult to extend these results to account for mobile crowders which are, of course, much more physically realistic. Preliminary investigations showed that both the starting theory, eq. 11, and the crowder proximity model, eq. 24, overestimate reaction rates even at very low crowding densities when the crowders diffuse. It’s conceivable that this discrepancy arises from mobile crowders being more effective at reducing access to reactants than stationary crowders, making the third term in eq. 24 too small. However, this seems unlikely because mobility would presumably make crowders less effective at restricting access to reactants, rather than more effective. A more likely explanation is that the reaction acceleration factor, [1 −*ϕ*_*x*_(*ϕ*)]^−1^, which applies to all of the terms in eq. 24, is too large for mobile crowders. In other words, mobile crowders effectively remove less volume from the system than their actual excluded volume, perhaps because their motion constantly opens up new volume, even while removing other volume. As an extreme example, an immobile water molecule would reduce the system volume that is available to proteins, but normal diffusing water molecules do not. The same arguments apply to the other models.

Two primary goals of this work were to build a deep understanding of reactions in crowded spaces and to create a theory that enables one to accurately convert between reaction rates in intracellular environments and those measured in dilute solutions. The work described here does not achieve either goal in its entirety but is a significant step in the right direction. It is substantially better than a previous theory, shows what types of effects are likely to be important, and identifies topics for further study.

## 4. Methods

Fields of randomly located crowded spheres were generated with SmolCrowd version 2 [27, 28]. These spheres had inner portions with radius *σ*_*c*_ that did not overlap and outer portions with radius *σ*_*BC*_ that could overlap (typically, *σ*_*C*_ = 0.5 nm and *σ*_*BC*_ = 1 nm). SmolCrowd’s algorithm is that it adds randomly placed spheres to the system volume one at a time, rejecting trial locations that would produce inner-portion overlap with an existing sphere. After many sequential failed trials (equal to ratio of the system to sphere volumes), SmolCrowd removes a randomly chosen sphere, and then returns to trying to add more. It continues until the requested crowding fraction is achieved. It ignores the existing sphere distribution when generating trial locations and when removing spheres in order to preserve an unbiased random crowder distribution.

SmolCrowd computed the occupied volume fraction, *ϕ*, using eq. 6 and the excluded volume fraction, *ϕ*_*x*_(*ϕ*), by randomly choosing 10^5^ points within the simulation volume and determining what fraction of them were within *σ*_*BC*_ of a sphere center. In simulations that had a central emitter (e.g. Figure 5D), spheres were prevented from having their inner edges within 1 nm of the origin or their outer edges beyond 10 nm from the origin. Other simulations used periodic boundary conditions, for which SmolCrowd duplicated all spheres that overlapped system boundaries as needed.

Diffusion and reaction simulations were run with Smoldyn version 2.61 [27, 47] using a cubical system that was 50 nm on each side, typically with periodic boundary conditions. All of Smoldyn’s algorithms approach exactness in the limits of short time steps, so simulations were made accurate by reducing time steps until simulation results stopped changing, and also did not change with a further 5-fold reduction in time step lengths; time steps ranged from 0.0002 µs to 0.005 µs. Simulations ran until all A molecules had reacted or for 10 µs of simulated time, whichever came first. Smoldyn represented the crowders generated by SmolCrowd as immobile reflective spherical surfaces; the most crowded simulations used about 63,000 crowders. Simulations typically started with 1000 each of A, B, and tracer molecules, each represented as simple points. Each diffused at 10 µm^2^/s, which is a typical intracellular diffusion coefficient [7]. The tracers did not interact with other molecules but were included to measure diffusion coefficients, which was done by quantifying their mean square displacements, ⟨*x*^2^⟩, and computing the diffusion coefficient from *D* = ⟨*x*^2^⟩/6Δ*t* where Δ*t* is the simulation time. Mean square displacements generally increased linearly over time, showing normal diffusion, although some anomalous subdiffusion [57, 58, 59] was observed for excluded volume fractions of 0.9 and above. The fact that diffusion was predominantly normal is consistent with the use of monodisperse crowders and simulations that ran long enough that most tracers diffused tens of times farther than the crowder radii.

Simulated reactions used the formula A+B → B, which simplified analysis by maintaining a constant B concentration. A molecules did not interact with other A molecules, or B molecules with other B molecules, including through excluded volume interactions, which simplified simulations and meant that excluded volume arose only from crowders. Reactions were interpreted using a model that I call the two-radius Smoluchowski model, in which reactants nominally interact when they are within a contact radius of each other, *σ*_*AB*_, but don’t actually react until they are at a smaller radius, *σ*_*T*_, where they always react. The reaction rate constant in this model is 4*πσ*_*T*_*D*, which is smaller than the diffusion-limited rate constant by a factor of *σ*_*T*_/*σ*_*AB*_, so the diffusion-limited fraction value in this model is *χ* = *σ*_*T*_/*σ*_*AB*_. This model is identical to the Collins and Kimball model for all radii that are larger than the contact radius. Because Smoldyn simulations use finite time steps, it used a slightly larger reaction radius than *σ*_*T*_ in order to produce the desired reaction rate constant, as described in ref. [26]. Steady-state rate constants in the crowded systems were quantified by fitting the number of A molecules in the system over time with an exponential decay function for reactions that were nearly activation-limited, or by fitting the time-dependent numerical rate constant with a function with the form 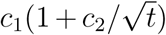 for reactions that were nearly diffusion-limited, where *c*_1_ and *c*_2_ are fitting parameters and *t* is the time [24]; this function has the same form as the time-dependent Smoluchowski reaction rate constant [31]. In all cases, these functions fit the data closely.

## 5. Acknowledgements

I thank Otto Berg, Ethan Hunt, Sam Isaacson, Satya Arjunan, and Hartmut Kuthan for helpful discussions. I also thank anonymous reviewers for helpful comments.

